# Transcriptomic analysis of the early strobilar development of *Echinococcus granulosus*

**DOI:** 10.1101/271767

**Authors:** João Antonio Debarba, Karina Mariante Monteiro, Alexandra Lehmkuhl Gerber, Ana Tereza Ribeiro de Vasconcelos, Arnaldo Zaha

## Abstract

**Background:** *Echinococcus granulosus* has a complex life cycle involving two mammalian hosts. The transition from one host to another is accompanied by changes in gene expression, and the transcriptional events that underlie these processes have not yet been fully characterized.

**Results:** In this study, RNA-seq is used to compare the transcription profiles of four time samples of *E. granulosus* protoscoleces *in vitro* induced to strobilar development. We identified 818 differentially expressed genes, which were divided into eight expression clusters formed over the entire 24 hours time course and indicated different transcriptional patterns. An enrichment of gene transcripts with molecular functions of signal transduction, enzymes and protein modifications was observed with progression of development.

**Conclusion:** This transcriptomic study provides insight for understanding the complex life cycle of *E. granulosus* and contributes for searching for the key genes correlating with the strobilar development, providing interesting hints for further studies.

## Background

Echinococcosis is a zoonotic parasitic infection caused by tapeworms of the genus *Echinococcus* and considered as one of the 17 neglected tropical diseases prioritized by the World Health Organization [1,2]. The two most important forms of the disease are cystic echinococcosis (hydatidosis) and alveolar echinococcosis, caused by infection with *Echinococcus granulosus* and *Echinococcus multilocularis*, respectively. *Echinococcus* has a two-host life cycle with the larval stage growing in the tissues of an intermediate host (large variety of non-carnivorous species) and the adult stage living in the intestine of a definitive host (few species of carnivores) [3,4].

*E. granulosus* larvae, also called metacestode, is a fluid-filled vesicular cyst containing many infectious protoscoleces, the pre-adult forms of the parasite. *In vitro* studies demonstrated that protoscoleces have an unusual potential of being able to differentiate in two different directions, depending on the environmental conditions provided. In the intermediate host, upon hydatid cyst rupture, protoscoleces released in the body cavity differentiate in a cystic direction (secondary hydatid cysts). In contrast, protoscoleces ingested by a dog and exposed to the gut environment, sexually differentiate in a strobilar direction to form fully developed, segmented adult tapeworms [4–7].

The strobilar development is directly influenced by the host-parasite relationship and configure a key point for the life cycle. Initially, the protoscolex remains quiescent and invaginated within the cyst until it receives the correct stimuli for the strobilation upon ingestion by a definitive host [7,8]. The nature of these stimuli is not fully known, but it is believed that chewing and the proteolytic enzymes like pepsin have a considerable role in this process, as well as temperature and the presence of bile salts. In addition, histological studies have already demonstrated that parasite binding or contact with a substrate similar to that found on the surface of the canine gut also constitutes an important stimulus for strobilation [9–11]. These observations resulted in the elaboration of strategies for *in vitro* culture of *Echinococcus* protoscoleces in order to provide the necessary physiological conditions for parasite strobilar development (summarized in Smyth, 1990).

In the molecular aspects, genome and transcriptome studies for *E. granulosus* [13–15] and *E. multilocularis* [14,16] identified differentially expressed genes between parasite life stages. Furthermore, considerable losses and gains of genes that may be associated with adaptations to parasitism were identified. In *E. granulosus*, crucial genes or even entire pathways of *de novo* synthesis of fatty acids, cholesterol, pyrimidines, purines and most amino acids are absent. Thus, *E. granulosus* relies on the host for obtaining these nutrients. Specifically in relation to strobilar development, it is known that bile acids have a crucial role in the differentiation of protoscoleces into adult worms [12], involving the expression of bile acid receptors and transporters to stimulate the pathways [14]. However, the association between these molecular events and the gradual morphological changes during the strobilar development of *E. granulosus* remain essentially unknown.

In an attempt to finding genes and molecular pathways involved with the gradual phenotypic changes triggered by strobilation stimuli in *E. granulosus*, we reported here the transcriptomic profile for the first 24 hours after protoscolex *in vitro* induction to adult development. We performed a comparative analysis of the transcriptomes from four different time-point samples during protoscolex strobilar induction and provide an overview of the early parasite developmental processes. Our data constitute the basis of future studies aimed at investigating the strobilar development and the host-parasite relationships that could be applied in the improvement of new control strategies for echinococcosis.

## Methods

### Parasite material and in vitro cultivation

*E. granulosus* protoscoleces (G1 genotype) were aseptically collected from a naturally infected liver of cattle routinely slaughtered at a commercial abattoir (São Leopoldo, RS, Brazil). The viability of protoscoleces was determined by trypan blue exclusion test and confirmed based on their motility characteristics under light microscopy [17]. Protoscoleces were washed 3 times with PBS, pH 7.4 (Sample 1 – PBS) and genotyped by one-step PCR [18]. PSCs were cultured *in vitro* as previously described [19]. Briefly, protoscoleces were incubated for 15 min with pepsin (2 mg/mL), pH 2.0 (Sample 2 – PEP), washed with PBS and transferred to a biphasic medium contained taurocholate for 12 (Sample 3 – 12 h) or 24 h (Sample 4 – 24 h). Additionally, protoscoleces were maintained in culture to confirm the characteristic morphological changes of strobilar development, as previously described [19].

### RNA extraction

Total RNA from each parasite sample was extracted using Trizol reagent, according to manufacturer’s instructions, followed by treatment with RNase-free DNase I (Sigma) to remove DNA contaminants. The integrity of the extracted RNA was monitored using gel electrophoresis on a 1% agarose gel. RNA concentration was determined using Qubit (Molecular Probes).

### cDNA library construction and sequencing

For cDNA libraries, 4 μg of total RNA were used as start material. PBS, PEP, 12 h and 24 h sample libraries were constructed, without replicate, using the TruSeq Stranded mRNA LT Sample Preparation Kit (Illumina) according to manufacturer’s instructions. Library quality control was performed using the 2100 Bioanalyzer System with the Agilent High Sensitivity DNA Kit (Agilent). The libraries were individually quantified via qPCR using a KAPA Library Quantification Kits for Illumina platforms (KAPA Biosystems). They were pooled together in equal amounts and sequenced in a MiSeq Sequencing System (Illumina). Paired-end reads (2×75 bp) were obtained using a MiSeq Reagents Kit v3 (150 cycles.)

### Data analysis

FastQC v0.11.2 [20] was used for data set quality checking. Individual Illumina read files (fastq) were trimmed and filtered using Trimmomatic v0.36 [21]. Paired end Trimmomatic parameters used were: LEADING:10 TRAILING:10 SLIDINGWINDOW:30:20 MINLEN:30. Filtered reads were mapped to *E. granulosus* genome by using TopHat2 v2.1.0 [22]. The genome of *E. granulosus* (PRJEB121, Tsai et al., 2013) and annotation (version 2014-05) were retrieved from WormBase ParaSite database [23].

The reads mapped to each transcript were used to calculate normalized transcript abundance and to perform differential gene expression analysis in GFOLD v1.1.4 [24], a software package specifically designed for unreplicated RNA-seq data. Genes with |GFOLD value|>1 or |log2 (fold change)|>2 were considered to be differentially expressed.

Hierarchical cluster analysis and Pearson correlation coefficient was performed using the R Stats Package v3.4.0, corrplot package v0.77 and RStudio v1.0.143. The conserved functional domain structures (SUPERFAMILY, Wilson et al., 2009) of the identified differentially expressed (DE) genes were predicted using InterProScan 5.21-60.0 [26]. The eggNOG database v4.5.1 [27] was used to acquire the functional annotation for the DE genes.

## Results

The paired-end RNA-seq sequencing using Illumina technology was used in order to investigate the transcriptome of protoscoleces induced to strobilar development. The RNA-seq resulted in a total of 30,821,916 reads. The overall raw read mean quality score was high, with 98.4% of bases above Q30. After quality filtering, 30.8 million of paired-end reads (99.5%) were obtained and 71.7% of the reads were mapped to the *E. granulosus* genome with known gene annotations. Table 1 shows the summary of the sequencing results.

**Table 1.** Summary of the *E, granulosus* RNA-seq data.

| **Samples** | **Q30<sup>a</sup>** | **Raw reads** | **Filtered reads** | **Mapped reads** |
| --- | --- | --- | --- | --- |
| **PBS** | 0.984 | 6,994,764 | 6,965,344 (99.6%) | 4,909,354 (70.5%) |
| **PEP** | 0.983 | 7,348,062 | 7,309,726 (99.5%) | 5,090,978 (69.6%) |
| **12 h** | 0.985 | 8,308,408 | 8,271,920 (99.6%) | 6,103,570 (73.8%) |
| **24 h** | 0.982 | 8,315,180 | 8,274,926 (99.5%) | 6,085,915 (73.5%) |

<sup>a</sup>Q30: Phred Quality Score; probability of incorrect base call: 1 in 1000.
PBS: protoscoleces washed with PBS; PEP: protoscoleces washed with PBS + treatment with pepsin; 12 h: protoscoleces subjected to PBS + PEP + biphasic medium for 12 h; 24 h: protoscoleces subjected to PBS + PEP + biphasic medium for 24 h.

A total of 9376 different genes were present in our dataset. A total of 9019, 9029, 9051 and 9077 genes were found in the PBS, PEP, 12 h and 24 h samples, respectively. Most of the genes (8742 genes) were detected in the four samples analyzed, but 61, 72, 56 and 73 genes were exclusively detected in PBS, PEP, 12 h and 24 h samples, respectively (Figure 1). A complete list of identified genes is provided in Additional file 1.

**Figure 1.**
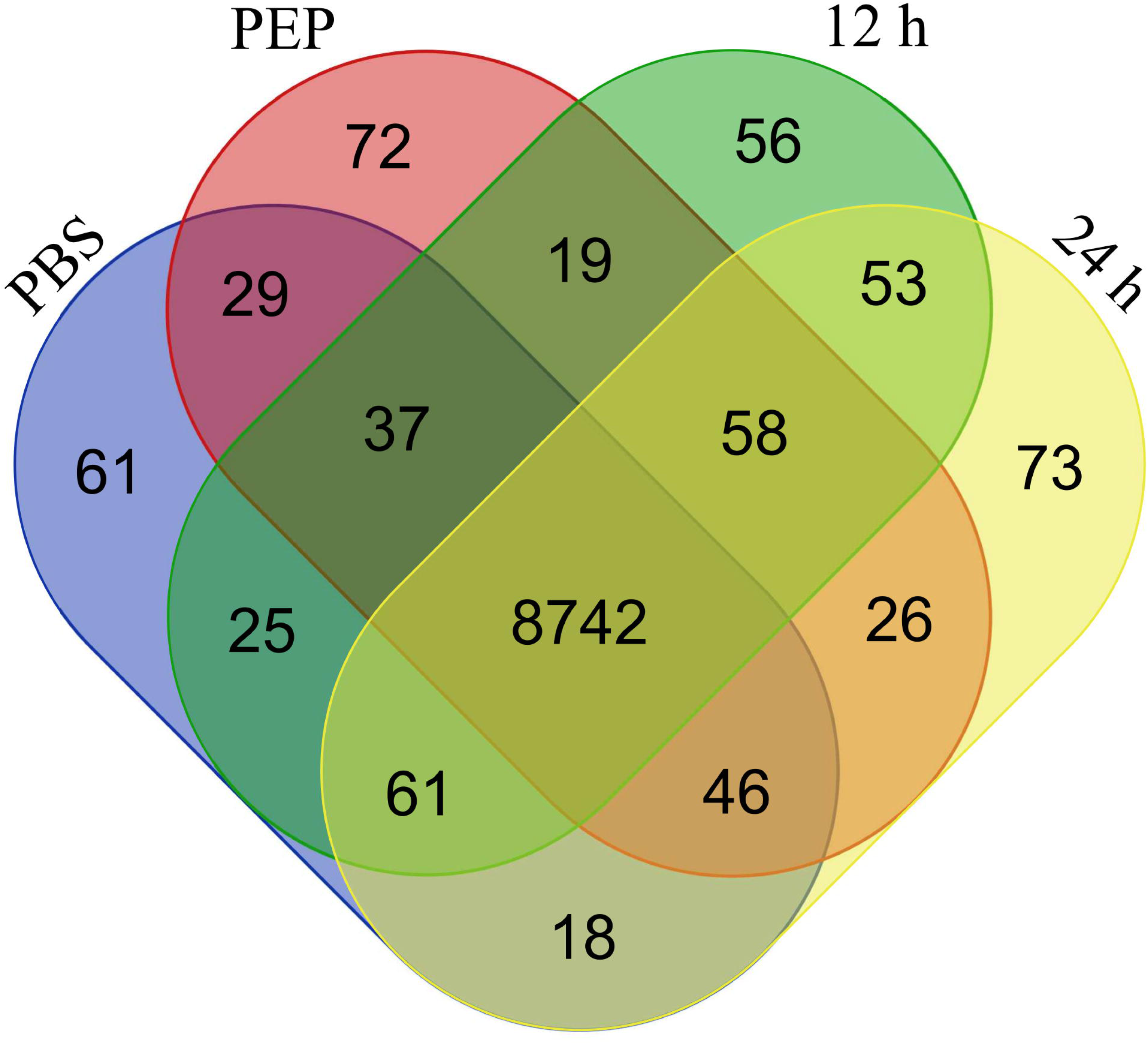
Venn diagram showing the distribution of detected genes between *E. granulosus* protoscolex samples. Genes with non-zero RPKM are represented and compared to show the number of genes with overlapping expression at four samples analyzed. The variation between protoscolex samples was calculated with the Pearson correlation. As shown in Figure 2, PBS and PEP samples show higher correlation coefficients than 12 and 24 h samples, indicating that there were little variation among them and a separation from the other samples.

**Figure 2.**
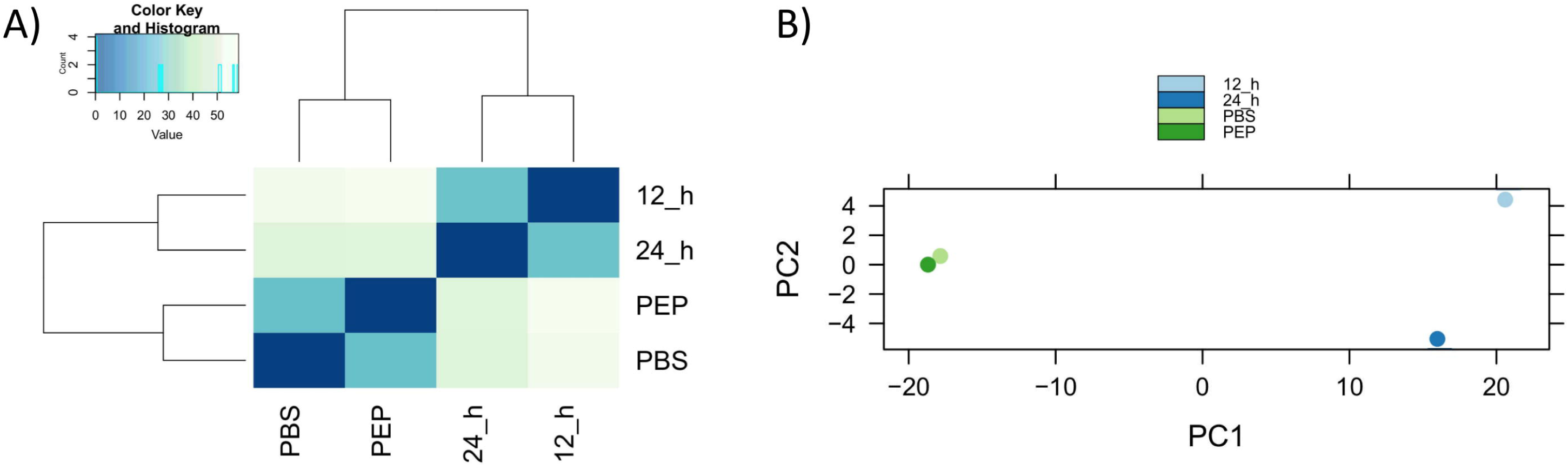
Correlation analysis of sequenced samples. (A) Pearson correlation coefficients between samples; (B) Principal Component Analysis.

We performed a pairwise comparison across different samples with GFOLD. A total of 818 genes were differentially expressed between any two samples (Additional file 2). The number of DE genes (up- and downregulated) in each 2×2 sample comparison is shown in Figure 3. A predominance of downregulated genes was detected in all pairwise comparisons. PEP *vs* 12 h showed the highest amount of DE genes (552 genes, 180 upregulated and 372 downregulated). Some of these data are shown in Table 2. These genes are related to growth and development, immunity, signal transduction, etc, and may be important for strobilation.

**Table 2.** Some of the genes that were differentially expressed among the samples analyzed of *E. granulosus*

| Gene_ID | Gene_Name | RPKM |  |  |  | Profile |  |  |  | Superfamily<sup>a</sup> | EggNOG<sup>b</sup> |
| --- | --- | --- | --- | --- | --- | --- | --- | --- | --- | --- | --- |
|  |  | ST | PEP | 12h | 24h | PBSx12 h | PBSx24 h | PEPx12 h | PEPx24 h |  |  |
| EgrG_000107200 | 2 amino 3 ketobutyrate coenzyme A ligase | 201,774 | 203,383 | 1583,37 | 1074,42 | Up | Up | Up | Up | RB | E |
| EgrG_000117000 | Ankyrin | 24,0945 | 17,9373 | 59,5489 | 24,153 | - | - | Up | - | RD | O |
| EgrG_001165500 | Ankyrin | 74,7373 | 67,9019 | 246,727 | 182,952 | Up | Up | Up | Up | RD | E |
| EgrG_000193700 | Annexin | 1095,57 | 1303,16 | 443,858 | 833,11 | Down | - | Down | - | IA | U |
| EgrG_000244000 | Annexin | 872,956 | 973,713 | 303,621 | 434,858 | Down | - | Down | Down | IA | U |
| EgrG_000330300 | Annexin | 953,654 | 973,326 | 467,045 | 427,206 | Down | Down | Down | Down | IA | U |
| EgrG_000124900 | Aquaporin 4 | 195,726 | 234,991 | 28,6044 | 41,8997 | Down | Down | Down | Down | RF | G |
| EgrG_000153200 | Aquaporin 9 AQP 9 small solute channel 1 | 6,07709 | 6,86419 | 325,38 | 156,091 | Up | Up | Up | Up | RF | G |
| EgrG_001190800 | Aquaporin 9 AQP 9 small solute channel 1 | 11,3987 | 8,44232 | 422,027 | 210,038 | Up | Up | Up | Up | RF | G |
| EgrG_000203700 | Choline transporter protein 2 | 72,4286 | 75,4026 | 30,3665 | 33,5379 | Down | - | Down | Down | - | P |
| EgrG_000127200 | Concentrative Na nucleoside cotransporter | 0,472568 | 1,20989 | 287,929 | 113,074 | Up | Up | Up | Up | - | P |
| EgrG_000342900 | Cysticercus cellulosae specific antigenic | 18289,6 | 20966,8 | 117,226 | 204,453 | Down | Down | Down | Down | - | - |
| EgrG_000261100 | Diagnostic antigen gp50 | 9,18014 | 6,89773 | 1,77943 | 2,37523 | Down | - | Down | - | - | - |
| EgrG_000564000 | Diagnostic antigen gp50 | 441,138 | 466,854 | 46,5439 | 81,7405 | Down | Down | Down | Down | - | - |
| EgrG_000520550 | Diagnostic antigen gp50 | 0,446094 | 0 | 8,28655 | 3,982 | Up | Up | Up | Up | - | - |
| EgrG_000304800 | Diagnostic antigen gp50 | 3,21728 | 2,40246 | 161,229 | 133,223 | Up | Up | Up | Up | - | - |
| EgrG_000324200 | Diagnostic antigen gp50 | 104,485 | 115,282 | 568,807 | 422,461 | Up | Up | Up | Up | - | - |
| EgrG_000566700 | Diagnostic antigen gp50 | 145,332 | 149,558 | 497,9 | 427,249 | Up | Up | Up | Up | - | - |
| EgrG_000071400 | Dynein light chain | 574,633 | 776,583 | 374,921 | 400,581 | - | - | Down | - | N | Z |
| EgrG_000941000 | Dynein light chain | 1522,5 | 2014,49 | 431,851 | 361,5 | Down | Down | Down | Down | N | Z |
| EgrG_000941100 | Dynein light chain | 2298,07 | 3266,25 | 512,348 | 697,2 | Down | Down | Down | Down | N | Z |
| EgrG_000940900 | Dynein light chain | 3620,31 | 4518,12 | 1492,48 | 1444,34 | Down | Down | Down | Down | N | Z |
| EgrG_000347100 | Elongation of very long chain fatty acids | 168,956 | 193,03 | 429,035 | 277,86 | Up | - | - | - | - | I |
| EgrG_000321300 | Elongation of very long chain fatty acids | 145,857 | 168,369 | 379,318 | 343,661 | Up | - | - | - | - | I |
| EgrG_000549800 | Fatty acid binding protein FABP2 | 1195,61 | 1125,07 | 497,465 | 454,409 | Down | Down | Down | Down | RF | I |
| EgrG_000549850 | Fatty acid binding protein FABP2 | 15557,9 | 17680,6 | 7030,25 | 5519,12 | Down | Down | Down | Down | RF | I |
| EgrG_000500800 | Forkhead box protein J3 | 6,74812 | 10,582 | 25,906 | 19,0748 | Up | - | - | - | LA | K |
| EgrG_000159700 | Forkhead box protein K1 | 56,2761 | 55,7012 | 16,4889 | 13,4864 | Down | Down | Down | Down | LA | K |
| EgrG_000179000 | Forkhead box protein O4 | 127,353 | 101,933 | 47,5055 | 44,8686 | Down | Down | Down | Down | LA | K |
| EgrG_000320300 | Forkhead box protein P4 | 84,3874 | 94,8522 | 42,8213 | 33,9381 | - | Down | Down | Down | LA | K |
| EgrG_000166300 | Forkhead box Q:D protein transcription | 57,4808 | 55,836 | 168,579 | 133,053 | Up | - | Up | - | LA | K |
| EgrG_000521000 | Heat shock protein 70 | 10,0821 | 6,12389 | 69,4092 | 45,7634 | Up | Up | Up | Up | RC | O |
| EgrG_000520950 | Heat shock protein 70 | 6,45234 | 6,57028 | 108,38 | 50,6147 | Up | Up | Up | Up | RC | O |
| EgrG_000554100 | Heat shock protein 70 | 37,0527 | 31,2413 | 324,553 | 280,569 | Up | Up | Up | Up | O | O |
| EgrG_000772900 | Homeobox protein ceh 26 | 9,33366 | 9,52123 | 2,16725 | 6,50904 | Down | - | Down | - | LA | K |
| EgrG_000040400 | Krueppel factor 10 | 559,331 | 589,885 | 127,123 | 114,181 | Down | Down | Down | Down | LA | K |
| EgrG_000608700 | Krueppel factor 5 | 317,941 | 424,699 | 35,579 | 28,4103 | Down | Down | Down | Down | LA | K |
| EgrG_000419100 | Kunitz protease inhibitor | 1520,32 | 1792,19 | 769,481 | 648,642 | - | Down | Down | Down | OA | - |
| EgrG_000445600 | Long chain fatty acid coenzyme A ligase 5 | 395,132 | 314,496 | 137,806 | 124,926 | Down | Down | Down | Down | RC | I |
| EgrG_000212700 | Major egg antigen p40 | 2923,9 | 3139,22 | 1186,33 | 848,683 | Down | Down | Down | Down | O | O |
| EgrG_000586900 | Oxalate:formate antiporter | 58,5919 | 46,314 | 22,5862 | 26,3801 | Down | - | - | - | P | G |
| EgrG_000586800 | Oxalate:formate antiporter | 9,41304 | 6,93566 | 56,9295 | 24,2628 | Up | - | Up | Up | P | G |
| EgrG_000017400 | Oxalate:formate antiporter | 186,078 | 190,454 | 28,2336 | 36,1629 | Down | Down | Down | Down | P | G |
| EgrG_000661800 | Oxalate:formate antiporter | 248,386 | 231,971 | 33,6004 | 48,2633 | Down | Down | Down | Down | P | G |
| EgrG_000204100 | P53 transcription factor DNA binding | 98,8238 | 81,228 | 39,9723 | 41,4273 | Down | Down | - | - | - | - |
| EgrG_000249100 | Phospholipase d1 | 0,193565 | 0,27876 | 19,9917 | 17,2063 | Up | Up | Up | Up | RC | I |
| EgrG_000204600 | Potassium voltage gated channel subfamily D | 6,35751 | 4,88303 | 0,944766 | 3,31039 | Down | - | - | - | P | P |
| EgrG_000197100 | Proto oncogene serine:threonine protein kinase | 147,414 | 116,318 | 48,2426 | 38,11 | Down | Down | Down | Down | OB | T |
| EgrG_001001900 | Pyruvate kinase | 0,569787 | 2,55289 | 5,50381 | 3,67332 | Up | Up | - | - | C | G |
| EgrG_001127400 | Sodium bile acid cotransporter | 0,767959 | 0,552983 | 5,13557 | 2,85629 | Up | - | Up | - | - | - |
| EgrG_001127500 | Sodium bile acid cotransporter | 13,8243 | 13,0956 | 76,5597 | 56,7643 | Up | Up | Up | Up | - | P |
| EgrG_000381100 | Tapeworm specific antigen B | 18,6673 | 6,72087 | 2,6007 | 1,73574 | Down | Down | - | - | - | - |
| EgrG_000381600 | Tapeworm specific antigen B | 59,6463 | 93,9159 | 252,619 | 231,605 | Up | Up | - | - | - | - |
| EgrG_000381400 | Tapeworm specific antigen B | 705,858 | 551,111 | 25,1401 | 58,1474 | Down | Down | Down | Down | - | - |
| EgrG_000381200 | Tapeworm specific antigen B | 2680,18 | 2204,6 | 171,536 | 310,395 | Down | Down | Down | Down | - | - |
| EgrG_000381700 | Tapeworm specific antigen B | 34,8619 | 42,5989 | 167,195 | 157,953 | Up | Up | Up | Up | - | - |
| EgrG_000381800 | Tapeworm specific antigen B | 140,34 | 217,926 | 2490,87 | 3299,76 | Up | Up | Up | Up | - | - |
| EgrG_000328400 | Tetraspanin | 35,7626 | 42,9192 | 59,7888 | 105,163 | - | Up | - | - | RE | S |
| EgrG_001077400 | Tetraspanin | 131,894 | 137,408 | 308,601 | 578,232 | - | Up | - | Up | RE | S |
| EgrG_000873600 | Universal stress protein | 2596,12 | 3003,99 | 501,033 | 436,72 | Down | Down | Down | Down | F | T |
| EgrG_000811100 | Universal stress protein | 13,3471 | 25,0464 | 224,943 | 152,537 | Up | Up | Up | Up | F | T |
| EgrG_001017500 | Zinc finger protein | 50,7582 | 65,6282 | 17,2905 | 17,7426 | Down | Down | Down | Down | LA | K |
| EgrG_000186000 | Zinc finger protein 45 | 13,5889 | 10,3948 | 1,88806 | 2,05454 | Down | Down | Down | Down | LA | K |
| EgrG_000711000 | Zinc finger transcription factor gli2 | 35,5351 | 41,3339 | 14,4482 | 11,1371 | Down | Down | Down | Down | LA | K |
| EgrG_000737300 | Zinc transporter foi | 18,7469 | 24,4141 | 59,7177 | 72,487 | Up | Up | - | Up | - | P |

<sup>a</sup>Superfamily code: C – energy; F - nucleotide metabolism and transport; IA – phospholipid metabolism and transport; LA – DNA-binding; N – cell motility; O – protein modification; OA – proteases; OB - kinases/phosphatases; P – ion metabolism and transport; RB – transferases; RC – other enzymes; RD – protein interaction; RE - immune response; RF – transport.
<sup>b</sup>EggNOG functional categories: E – amino acid transport and metabolism; G – carbohydrate transport and metabolism; I – lipid transport and metabolism; K – transcription; O – post-translational modification, protein turnover, and chaperones; P – inorganic ion transport and metabolism; S – function unknown; T – signal transduction mechanisms; U – intracellular trafficking, secretion, and vesicular transport; Z – cytoskeleton.

**Figure 3.**
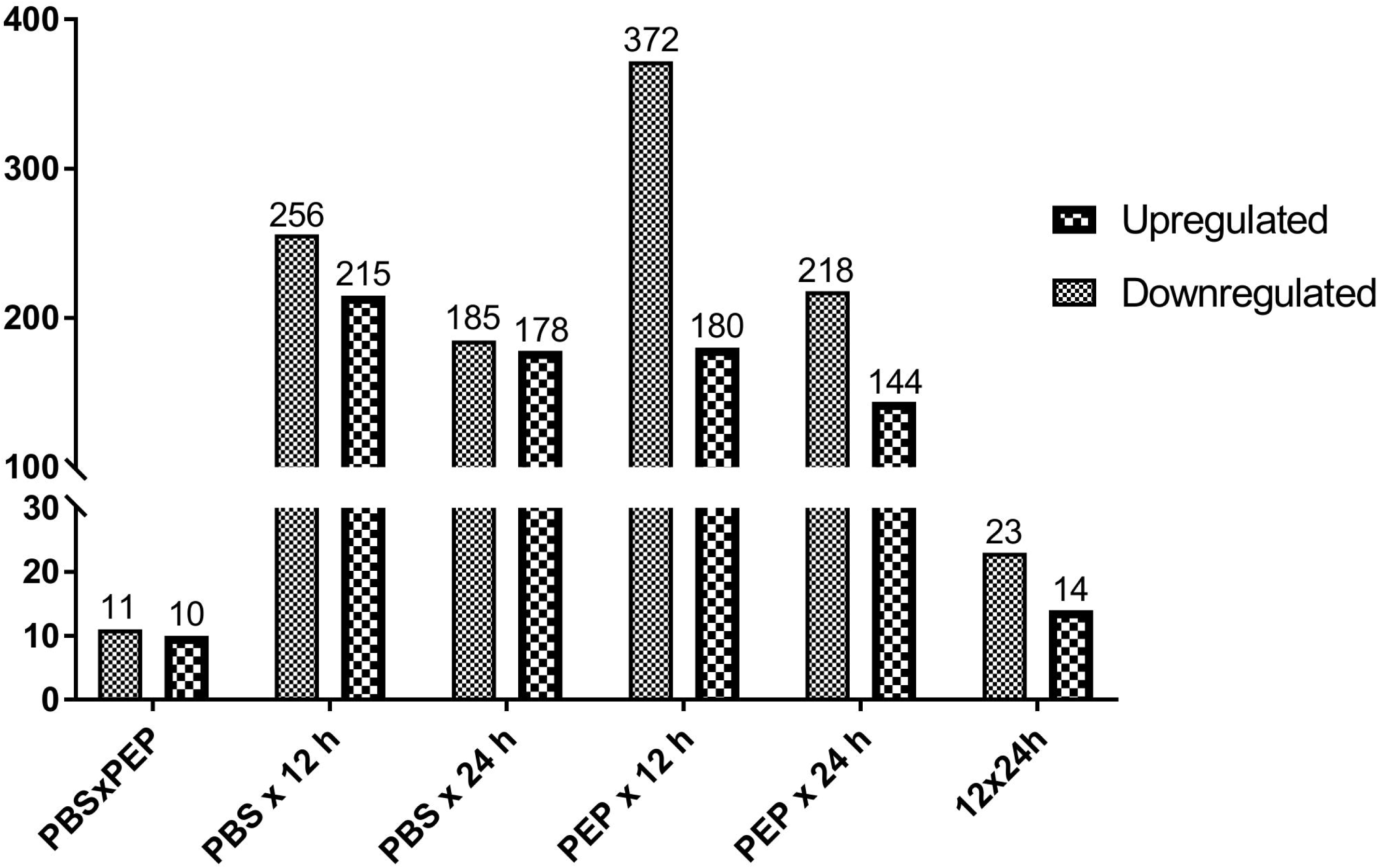
Differentially expressed genes. Up- and downregulated genes in each pairwise comparison are shown.

To better understanding of the dynamics of *E. granulosus* protoscolex gene expression, DE genes were clustered to search for similar patterns of expression between samples (Figure 4). Using the sum of the RPKMs values in the four samples (cutoff > 10), 744 DE genes were categorized into eight different clusters on account of their relative expression pattern (Additional file 2). We computed a relative expression for each gene by dividing its expression at each time point by the sum of gene expression for all time points. In this analysis, clusters 1 and 2, including 446 genes, showed decreasing expression profile patterns throughout the treatment for protoscolex strobilation, while 277 genes in clusters 6, 7 and 8 showed increasing expression profiling patterns.

**Figure 4.**
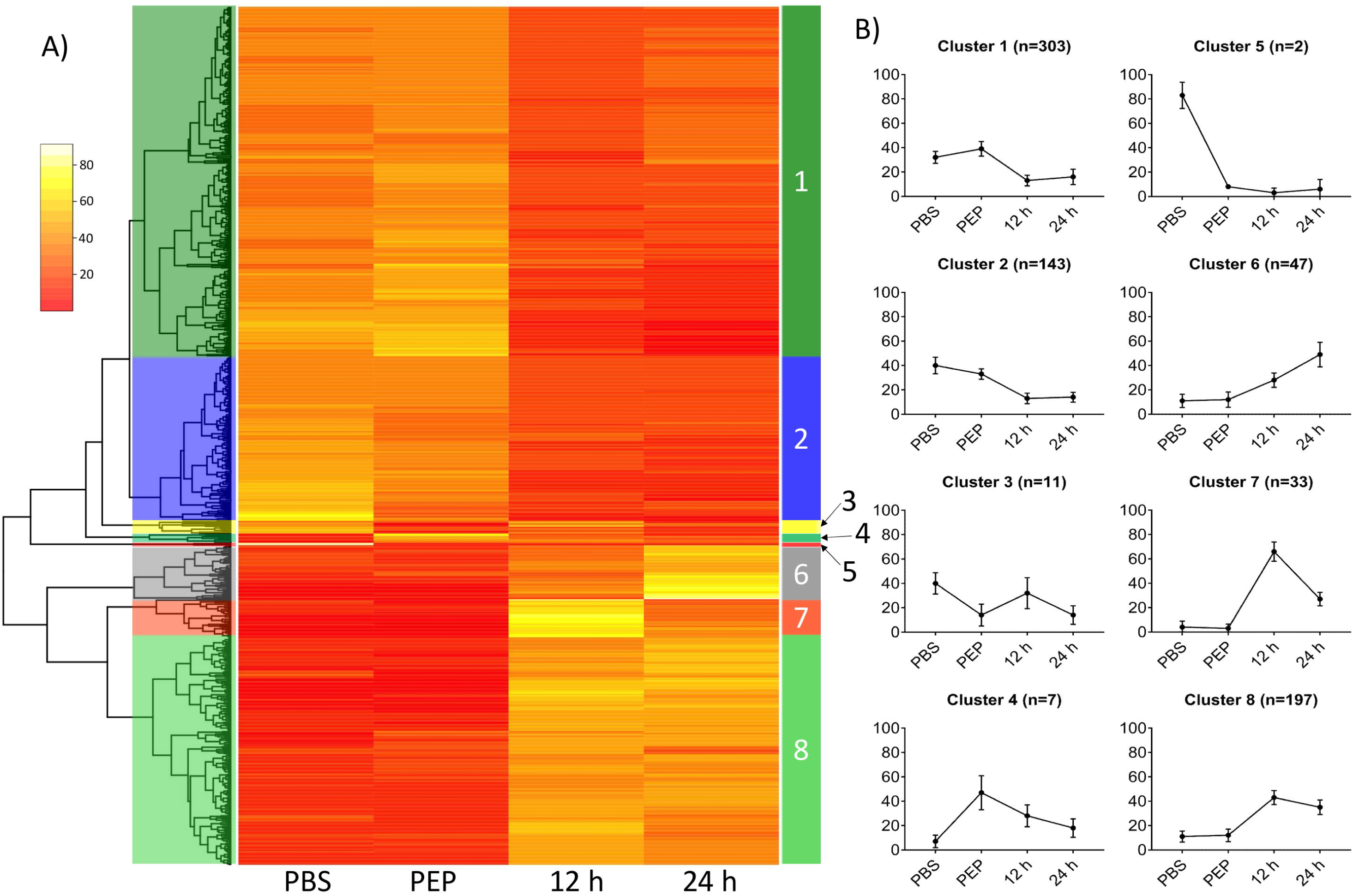
Heatmap plot of DE genes during *E. granulosus* strobilar development induction. (A) Relative expression of each gene is represented in the rows and categorized clusters; (B) Average expression (+/-standard deviation) for genes in each cluster reveals similar expression patterns.

Among the upregulated genes obtained from GFOLD analysis, we identified genes coding for 2-amino-3-ketobutyrate coenzyme A ligase, heat shock protein 70 (16 genes), sodium bile acid cotransporter (2 genes) and tetraspanin (4 genes). In contrast, among the downregulated genes, genes coding for annexin (6 genes), calcium binding protein (2 genes), dynein light chain (10 genes) and fatty acid binding protein FABP2 were identified.

A structural and functional annotation of the DE genes is summarized in Figure 5 (Additional file 2). The most representative domains found in the genes downregulated during early worm development (cluster 1and 2) were related to cell motility, cell adhesion, DNA-binding, translation and DNA replication/repair. Among the genes upregulated during early worm development (cluster 6, 7 and 8), the most representative functions were signal transduction, other enzymes and protein modification. By EggNOG analysis, the most significant functions found in downregulated genes were translation, ribosomal structure and biogenesis (J), intracellular trafficking, secretion, and vesicular transport (U) and cytoskeleton (Z). In the upregulated genes, the most relevant functions were inorganic ion transport and metabolism (P), nucleotide transport and metabolism (F) and amino acid transport and metabolism (E).

**Figure 5.**
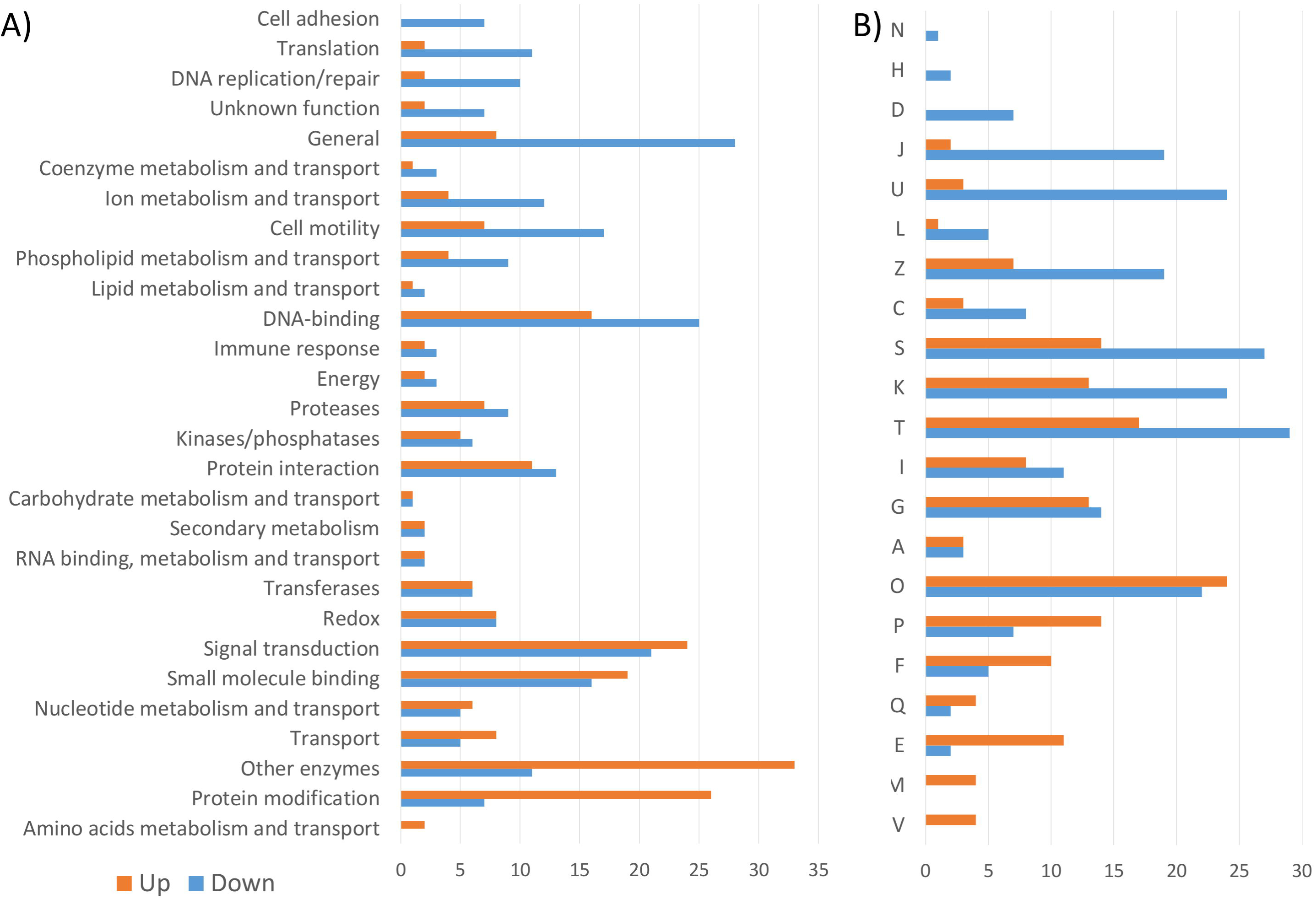
Comparative structural and functional analysis of upregulated (clusters 6, 7 and 8) and downregulated (clusters 1and 2) DE genes. (A) SUPERFAMILY predictions of the conserved domains shown by percentage; (B) EggNOG functional categories of DE genes shown by percentage. (A) RNA processing and modification; (C) Energy production and conversion; (D) Cell cycle control, cell division, chromosome partitioning; (E) Amino acid transport and metabolism; (F) Nucleotide transport and metabolism; (G) Carbohydrate transport and metabolism; (H) Coenzyme transport and metabolism; (I) Lipid transport and metabolism; (J) Translation, ribosomal structure and biogenesis; (K) Transcription; (L) Replication, recombination and repair; (M) Cell wall/membrane/envelope biogenesis; (N) Cell motility; (O) Post-translational modification, protein turnover, and chaperones; (P) Inorganic ion transport and metabolism; (Q) Secondary metabolites biosynthesis, transport, and catabolism; (S) Function unknown; (T) Signal transduction mechanisms;(U) Intracellular trafficking, secretion, and vesicular transport; (V) Defense mechanisms; (Z) Cytoskeleton.

Uncharacterized proteins represent 270 of total DE genes, with 68 annotated as expressed conserved proteins, 62 expressed proteins and 140 hypothetical proteins (Additional file 2). Of these, only 10 have conserved domains predicted by SUPERFAMILY and 13 have EggNOG functional categories (three of them present both classifications).

## Discussion

Important morphological and biochemical changes occur throughout the life cycle of parasitic organisms and are probably the result of regulated changes in gene expression in response to environmental stimuli such as host, temperature and pH changes [28–30]. These regulated responses contribute to the mechanisms by which parasites subvert host immune defenses and cause infection.

Previously, based on the classic works of Smyth and collaborators (summarized in Smyth, 1990), we have reported an *in vitro* culture of *E. granulosus* protoscolex strobilar development based on a biphasic medium containing the bile salt taurocholate [19]. In this work, we cultivate protoscoleces in biphasic medium for 12 or 24 hours. In addition, we used untreated protoscoleces washed with PBS (PBS) and protoscoleces treated with pepsin (PEP) to compare and to search for DE genes involved in the strobilar development.

Based on the Jacob-Monod model, a hypothetical but logical model was proposed to explain how the gene expression regulation can be involved in the *Echinococcus* development [7,31]. Although both the morphological characteristics of the strobilar development and the genome of *E. granulosus* are known, the correlation between these two informations and the model previously proposed is still poorly understood. In this work, we provide a transcriptional analysis of *E. granulosus* protoscoleces *in vitro* induced to strobilar development in attempt to finding genes involved in this process. By sample-to-sample correlation analysis, it was possible to observe that the most abundant transcripts share similarities between the different samples analyzed. PBS and PEP samples have a relative high-level expression of several identical gene transcripts. Furthermore, with the subsequent activation of the protoscoleces by pepsin and bile salts, mimicking the developmental transition in the definitive host, a change in the identity of highly expressed genes can be observed.

Among the most expressed genes found in our data, it is important to verify the presence of fatty acid binding protein (FABP) and the antigen B transcripts. Both of them have already been described among the highly *Echinococcus* expressed genes [14,32], which corroborates the validity of our data. The importance of these genes lies in the fact that cestodes are unable to synthesize fatty acids and cholesterol *de novo* (Tsai et al., 2013). They depend essentially on the sequestration and utilization of host lipids by proteins like FABP and the antigen B. In this study, FABP genes are downregulated during *E. granulosus* adult worm development. Antigen B has both up- and downregulated genes, in agreement with previous works [33,34].

Among the downregulated DE genes, we found several genes coding for dynein light chain, oxalate:formate antiporter and annexins. Dynein is a family of cytoskeletal motor proteins involved in intracellular motility of vesicles and organelles along microtubules and are associated with transforming growth factor (TGF)-β signaling [14,35]. Previous studies showed the expansion of this family in *E. granulosus* and schistosomes when compared to nematodes [13,14]. Oxalate:formate antiporter is a subfamily of the major facilitator transporter family, responsible for the transport of small solutes [36], but its function is not fully understood in parasites. Annexins, in contrast, are considered to play critical roles in parasite process related to the maintenance of cell integrity and modulation of the host immune responses [37]. Therefore, the decrease in the expression of the annexins may be related to the absence of contact with the host in the *in vitro* cultures.

On the other hand, we found ankyrin, tetraspanin, heat shock protein 70 (Hsp70) and sodium bile acid cotransporter among upregulated DE genes. Ankyrins are involved in functions such as cell-cycle regulation, transcriptional regulation, cytoskeleton interactions, signal transduction, development and intracellular trafficking [38,39]. In parasites, tetraspanins are involved in the coordination of signal transduction, cell proliferation, adhesion, migration, cell fusion, and host–parasite interactions [40]. In *E. granulosus*, tetraspanins were mostly present in the tegument and could contribute to the parasite nutrition [41]. The Hsp70 are part of the group of the expanded domain families in *E. granulosus* and may have important roles in protein folding and in protecting cells from stress [14]. The expression of Hsp70 may be particularly associated to the stressful conditions of strobilation induction, which involves an increase in protein synthesis [19,42]. In turn, sodium bile acid cotransporter is an integral membrane glycoprotein that, in humans, participate in the enterohepatic circulation of bile acids. Bile acids seem to play a key role in the differentiation of *Echinococcus* protoscoleces into adult worms, and the expression of bile acid receptors and transporters may be stimulated during strobilar development [12,15].

In our previous work, we identified proteins expressed by *E. granulosus* protoscoleces upon the induction of strobilar development [19]. These proteins were related to the cytoskeleton, energy metabolism and cellular communication. Specifically, the 2-amino-3-ketobutyrate coenzyme A ligase, classified here as an upregulated DE gene, had an increased expression in the presence of strobilation stimuli.

When we analyzed the molecular function of DE genes, we also found differences between clusters. In cluster 1, which presents a downregulated expression pattern, we observed the presence of more basal functions, such as translation (e.g., Eukaryotic translation initiation factor 5a), DNA replication and cell motility. In contrast, the upregulated DE genes of cluster 8 are related to specialized functions like signal transduction (e.g., tyrosine protein kinase and G protein coupled receptor), enzymes (e.g., hexokinase and phospholipase) and protein modifications (e.g., Hsp70), which might correlate with the increased morphological complexity of the adult tapeworm compared to the metacestode. It is important to note that an expressive number of genes (270 in DE genes; 2976 in total) are not characterized, which difficult more accurate analyses. This is the case, for example, of the hypothetical protein EgrG_000335800, which is one of the most expressed genes in the four conditions analyzed.

## Conclusion

In this study, we have conducted RNA-Seq analysis of the protoscoleces and early strobilar stages of *E. granulosus*. More importantly, this work provides information about DE genes in key intermediate stages, providing novel information about *E. granulosus* strobilar development. In summary, we provide here significant data that can be used to explore basic questions on the biology and evolution of cestodes, including the study of development and the host-parasite relationship.

## Additional file

Additional file 1: List of identified genes. (XLSX 2000 kb)

Additional file 2: List of differentially expressed genes. (XLSX 98 kb)

## Abbreviations

DE: differentially expressed
FABP: fatty acid binding protein
HSP: heat shock protein
RPKM: reads per kilobase million

## Availability of data and materials

All sequence data (raw Illumina reads) are available on the NCBI Sequence Read Archive (SRA) database under the accession ID SRP131874.

## Author’s contributions

JAD, KMM and AZ conceived and designed the experiments. JAD, SME, and ALG performed the experiments. JAD, KMM and AZ analyzed the data. JAD, ALG, ATRV and AZ contributed to the writing of the manuscript. All authors read and approved the final manuscript.

## Ethics approval and consent to participate

Not applicable.

## Consent for publication

Not applicable.

## References

1. da Silva AM. Human echinococcosis: A neglected disease. Gastroenterol. Res. Pract. [Internet]. 2010;2010. Available from: http://www.ncbi.nlm.nih.gov/pubmed/20862339

2. WHO. Investing to overcome the global impact of neglected tropical diseases [Internet]. Geneva; 2015. Available from: http://www.who.int/neglected_diseases

3. Eckert J, Deplazes P. Biological, Epidemiological, and Clinical Aspects of Echinococcosis, a Zoonosis of Increasing Concern. Clin. Microbiol. Rev. 2004;17:107–35.

4. Thompson RCA, Jenkins DJ. Echinococcus as a model system: Biology and epidemiology. Int. J. Parasitol. 2014;44:865–77.

5. Cucher M, Prada L, Mourglia-Ettlin G, Dematteis S, Camicia F, Asurmendi S, et al. Identification of Echinococcus granulosus microRNAs and their expression in different life cycle stages and parasite genotypes. Int. J. Parasitol. 2011;41:439–48.

6. Thompson RCA. Biology and systematics of echinococcus. Echinococcus hydatid Dis. Wallingford, Oxfordshire, UK: CAB International; 1995. p. 1–50.

7. Smyth JD. Parasites as biological models. Parasitology. Cambridge University Press; 1969;59:73.

8. Smyth JD, Miller HJ, Howkins AB. Further analysis of the factors controlling strobilization, differentiation, and maturation of Echinococcus granulosus in vitro. Exp. Parasitol. 1967;21:31–41.

9. Smyth J, Howkins A, Barton M. Factors controlling the differentiation of the hydatid organism, Echinococcus granulosus, into cystic or strobilar stages in vitro. Nature. Nature Publishing Group; 1966;211:1374–7.

10. Morseth DJ. Fine structure of the hydatid cyst and protoscolex of Echinococcus granulosus. J Parasitol. 1967;53:312–25.

11. Smyth JD, Gemmell M, Smyth MM. Establishment. of Echinococcus granuiosus in the intestine of normal and vaccinated dogs. Singh KS, Tandan BK, editors. Izatnagar, U.P.: Indian Veterinary Research Institute.; 1970.

12. Smyth JD. Cestoda. In: Smyth JD, editor. Vitr. Cultiv. Parasit. Helminthes. London: CRC press; 1990. p. 123–37.

13. Parkinson J, Wasmuth JD, Salinas G, Bizarro C V., Sanford C, Berriman M, et al. A Transcriptomic Analysis of Echinococcus granulosus Larval Stages: Implications for Parasite Biology and Host Adaptation. PLoS Negl. Trop. Dis. [Internet]. 2012;6:e1897. Available from: http://www.ncbi.nlm.nih.gov/pubmed/23209850

14. Tsai IJ, Zarowiecki M, Holroyd N, Garciarrubio A, Sanchez-Flores A, Brooks KL, et al. The genomes of four tapeworm species reveal adaptations to parasitism. Nature [Internet]. 2013;496:57–63. Available from: http://www.ncbi.nlm.nih.gov/pubmed/23485966%5 http://www.pubmedcentral.nih.gov/articlerender.fcgi?artid=PMC3964345

15. Zheng H, Zhang W, Zhang L, Zhang Z, Li J, Lu G, et al. The genome of the hydatid tapeworm Echinococcus granulosus. Nat. Genet. [Internet]. 2013;45:1168–75. Available from: http://dx.doi.org/10.1038/ng.2757

16. Huang F, Dang Z, Suzuki Y, Horiuchi T, Yagi K, Kouguchi H, et al. Analysis on Gene Expression Profile in Oncospheres and Early Stage Metacestodes from Echinococcus multilocularis. Zhang W, editor. PLoS Negl. Trop. Dis. [Internet]. Springer; 2016 [cited 2017 Apr 20];10:e0004634. Available from: http://dx.plos.org/10.1371/journal.pntd.0004634

17. Zhang R, Aji T, Shao Y, Jiang T, Yang L, Lv W, et al. Nanosecond pulsed electric field (nsPEF) disrupts the structure and metabolism of human Echinococcus granulosus protoscolex in vitro with a dose effect. Parasitol. Res. Springer Berlin Heidelberg; 2017;116:1345–51.

18. Balbinotti H, Santos GB, Badaraco J, Arend AC, Graichen DAS, Haag K are. L, et al. Echinococcus ortleppi (G5) and Echinococcus granulosus sensu stricto (G1) loads in cattle from Southern Brazil. Vet. Parasitol. 2012;188:255–60.

19. Debarba JA, Monteiro KM, Moura H, Barr JR, Ferreira HB, Zaha A. Identification of Newly Synthesized Proteins by Echinococcus granulosus Protoscoleces upon Induction of Strobilation. PLoS Negl. Trop. Dis. Public Library of Science; 2015; 9.

20. Andrews S. FastQC A Quality Control tool for High Throughput Sequence Data [Internet]. 2007. Available from: http://www.bioinformatics.babraham.ac.uk/projects/fastqc/

21. Bolger AM, Lohse M, Usadel B. Trimmomatic: A flexible trimmer for Illumina sequence data. Bioinformatics. 2014;30:2114–20.

22. Kim D, Pertea G, Trapnell C, Pimentel H, Kelley R, Salzberg SL. TopHat2: accurate alignment of transcriptomes in the presence of insertions, deletions and gene fusions. Genome Biol. BioMed Central; 2013;14:R36.

23. Howe KL, Bolt BJ, Shafie M, Kersey P, Berriman M. WormBase ParaSite - a comprehensive resource for helminth genomics. Mol. Biochem. Parasitol. 2017;215:2–10.

24. Feng J, Meyer CA, Wang Q, Liu JS, Liu XS, Zhang Y. GFOLD: A generalized fold change for ranking differentially expressed genes from RNA-seq data. Bioinformatics. Oxford University Press; 2012;28:2782–8.

25. Wilson D, Pethica R, Zhou Y, Talbot C, Vogel C, Madera M, et al. SUPERFAMILY--sophisticated comparative genomics, data mining, visualization and phylogeny. Nucleic Acids Res. Oxford University Press; 2009;37:D380–6.

26. Jones P, Binns D, Chang HY, Fraser M, Li W, McAnulla C, et al. InterProScan 5: Genome–scale protein function classification. Bioinformatics. Oxford University Press; 2014;30:1236–40.

27. Huerta-Cepas J, Szklarczyk D, Forslund K, Cook H, Heller D, Walter MC, et al. eggNOG 4.5: a hierarchical orthology framework with improved functional annotations for eukaryotic, prokaryotic and viral sequences. Nucleic Acids Res. Springer Science & Business Media, NY; 2016;44:D286–93.

28. Kramer S. Developmental regulation of gene expression in the absence of transcriptional control: The case of kinetoplastids. Mol. Biochem. Parasitol. 2012;181:61–72.

29. Thorson RE. Environmental Stimuli and the Responses of Parasitic Helminths. Bioscience. 1969;19:126–30.

30. Haile S, Papadopoulou B. Developmental regulation of gene expression in trypanosomatid parasitic protozoa. Curr. Opin. Microbiol. 2007;10:569–77.

31. Thompson RCA, Lymbery AJ. Let’s not forget the thinkers. Trends Parasitol. [Internet]. 2013 [cited 2017 Mar 7];29:581–4. Available from: http://linkinghub.elsevier.com/retrieve/pii/S1471492213001670

32. Obal G, Ramos AL, Silva V, Lima A, Batthyany C, Bessio MI, et al. Characterisation of the Native Lipid Moiety of Echinococcus granulosus Antigen B. Dalton JP, editor. PLoS Negl. Trop. Dis. Elsevier; 2012;6:e1642.

33. Espínola SM, Ferreira HB, Zaha A. Validation of suitable reference genes for expression normalization in Echinococcus spp. larval stages. PLoS One. 2014;9:e102228.

34. Mamuti W, Sako Y, Xiao N, Nakaya K, Nakao M, Yamasaki H, et al. Echinococcus multilocularis: Developmental stage-specific expression of Antigen B 8-kDa-subunits. Exp. Parasitol. 2006;113:75–82.

35. Roberts AJ, Kon T, Knight PJ, Sutoh K, Burgess SA. Functions and mechanics of dynein motor proteins. Nat. Rev. Mol. Cell Biol. Europe PMC Funders; 2013;14:713–26.

36. Pao SS, Paulsen IT, Saier MH. Major facilitator superfamily. Microbiol. Mol. Biol. Rev. American Society for Microbiology; 1998;62:1–34.

37. Cantacessi C, Seddon JM, Miller TL, Leow CY, Thomas L, Mason L, et al. A genome-wide analysis of annexins from parasitic organisms and their vectors. Sci. Rep. 2013;3:2893.

38. Siozios S, Ioannidis P, Klasson L, Andersson SGE, Braig HR, Bourtzis K. The diversity and evolution of Wolbachia ankyrin repeat domain genes. PLoS One. Public Library of Science; 2013;8:e55390.

39. Stankewich MC, Moeckel GW, Ji L, Ardito T, Morrow JS. Isoforms of Spectrin and Ankyrin Reflect the Functional Topography of the Mouse Kidney. PLoS One. Public Library of Science; 2016;11:e0142687.

40. Silva LL, Marcet-Houben M, Nahum LA, Zerlotini A, Gabaldón T, Oliveira G. The Schistosoma mansoni phylome: using evolutionary genomics to gain insight into a parasite’s biology. BMC Genomics. BioMed Central; 2012;13:617.

41. Hu D, Song X, Xie Y, Zhong X, Wang N, Zheng Y, et al. Molecular insights into a tetraspanin in the hydatid tapeworm Echinococcus granulosus. Parasit. Vectors. BioMed Central; 2015;8:311.

42. Cui SJ, Xu LL, Zhang T, Xu M, Yao J, Fang CY, et al. Proteomic characterization of larval and adult developmental stages in Echinococcus granulosus reveals novel insight into host-parasite interactions. J. Proteomics [Internet]. 2013;84:158–75. Available from: http://www.ncbi.nlm.nih.gov/pubmed/23603110

